# RECOMBINATION HOTSPOTS IN SOYBEAN [*GLYCINE MAX* (L.) MERR.]

**DOI:** 10.1101/2022.09.15.508170

**Authors:** Samantha McConaughy, Keenan Amundsen, Qijian Song, Vince Pantalone, David Hyten

## Abstract

Recombination allows for the exchange of genetic material between two parents which plant breeders exploit to make new and improved cultivars. This recombination is not distributed evenly across the chromosome. In crops, recombination mostly occurs in euchromatic regions of the genome and even then, recombination is focused into clusters of crossovers termed recombination hotspots. Understanding the distribution of these hotspots along with the sequence motifs associated with them may lead to methods that enable breeders to better exploit recombination in breeding. To map recombination hotspots and identify sequence motifs associated with hotspots in soybean [Glycine max (L.) Merr.], two bi-parental recombinant inbred lines (RILs) populations were genotyped with 50,000 SNP markers using the SoySNP50k Illumina Infinium assay. A total of 451 recombination hotspots were identified in the two populations. Despite being half-sib populations, only 18 hotspots were in common between the two populations. While pericentromeric regions did exhibit extreme suppression of recombination, twenty-seven percent of the detected hotspots were located in the pericentromic regions of the chromosomes. Two genomic motifs associated with hotspots are similar to human, dog, rice, wheat, drosophila, and arabidopsis. These motifs were a CCN repeat motif and a poly-A motif. Genomic regions spanning other hotspots were significantly enriched with the tourist family of mini-inverted-repeat transposable elements (MITEs) that resides in less than 0.34% of the soybean genome. The characterization of recombination hotspots in these two large soybean bi-parental populations demonstrates that hotspots do occur throughout the soybean genome and are enriched for specific motifs but their locations may not be conserved between different populations.

## INTRODUCTION

Recombination is a fundamental process that drives the rearrangement of alleles resulting in new combinations of those alleles. This can lead to favorable allelic combinations or break up undesirable ones, such as decoupling linked deleterious alleles (Barton and Charlesworth, 1998). However, recombination is a highly regulated process that involves DNA double stranded breaks, resulting in crossover or non-crossover events (Baudat *et al*., 2013) Crossovers, at least one per chromosome, are required for proper chromosome segregation (Mercier *et al*., 2015a).

The distribution, associated motifs, strength, and size of recombination vary across all eukaryotes, potentially implying a lack of unifying characteristics (Choi and Henderson, 2015; Choulet *et al*., 2014; Darrier *et al*., 2017; Marand *et al*., 2017; Mercier *et al*., 2015b; Saintenac *et al*., 2009; Tenaillon *et al*., 2002).

While the genomic landscape of recombination varies across plant species, it transpires in a small percentage of the genome. Recombination rates are the highest in gene-rich euchromatic regions and lowest in repeat-rich heterochromatic regions (Choulet *et al*., 2014; Gore *et al*., 2009; Li *et al*., 2015; Liu *et al*., 2009; Paterson *et al*., 2009; Rodgers-Melnick *et al*., 2015; Song *et al*., 2016; Wei *et al*., 2009). In soybean [*Glycine max* (L.) Merr.], 93% of recombination occurs in the euchromatic DNA which accounts for 43% of the genome (Schmutz *et al*., 2010). Recombination rates across the euchromatic regions in the soybean genome average one centimorgan (cM) per 197 kilobase (kb) while in pericentromeric regions the average increases to one cM per 3.5 megabase (Mb) (Schmutz *et al*., 2010).

Due to the non-random pattern of recombination, the relationship between the physical distance and genetic map distance results in it not being a one-to-one ratio across the chromosome. Crossovers are clustered into small physical distances that are defined as recombination hotspot regions. These recombination hotspots are often flanked by regions of DNA that have lower than expected recombination rates called recombination cold spots. In plants, hotspot regions have been discovered to vary in size from 500 to 23,000 base pairs (bp), whereas cold spots can range from 5,000 to millions of bp in length (Drouaud *et al*., 2013b; Fu *et al*., 2002; Patterson *et al*., 1995; Saintenac *et al*., 2009; Yao and Schnable, 2005; Yelina *et al*., 2012). The resolution to identify recombination hotspots and cold spots is often limited by marker density and population size.

While the size of recombination hotspots and cold spots are an important attribute, the distribution of recombination influences the probability of combining new allelic combinations and introgressing diversity in crops. In maize and wheat, recombination hotspots are located in sub-telomeric regions, whereas cold spots are concentrated near centromeres, telomeres, and interstitial regions (Lambing et al. 2017). In contrast, recombination hotspots in Arabidopsis are not concentrated in specific regions, but are dispersed throughout the chromosome, except in the centromere (Salome *et al*., 2012). Lambing *et al*. (2017) hypothesized the difference between Arabidopsis, maize, and wheat hotspot locations may be due to transposable elements.

While the distribution of hotspots varies by species and genome locations, the relationship between transposable elements along with sequence motif may help explain this difference. Several experiments have identified closely associated repeat motifs to recombination hotspots. The first discovery was in humans with a CCN-like motif bound by PRDM9 (PR/SET Domain 9) a zinc finger protein with histone methyltransferace activity (Myers *et al*., 2008). The 13-mer motif, CCNCCNTNNCCNC, was reported in 40% of human hotspots (Myers *et al*., 2008). In plants, three common motifs have been found in associations with hotspots including a CCN-repeat, CTT-repeat, and poly-A stretch motif (Choi *et al*., 2013; Shilo *et al*., 2015; Wijnker and de Jong, 2008). The annotation of the CCN-like motif discovered in wheat is related with the terminal inverted repeat (TIR)-Mariner sequence. These motifs are suggested to be involved in chromatin structure within promoter regions due to the location near transcription start sites and the similarity to PRDM9 protein (Darrier *et al*., 2017). Since plants do not have a known homolog for PRDM9, the DNA transposons from the Mariner family along with the associated motifs could be functionally related to recombination due to their high association with recombination hotspots (Darrier *et al*., 2017) These experiments provide insight into a potential relationship between transposable elements and recombination.

The development of the high-density genotyping array, SoySNP50K, has made it possible to conduct recombination hotspot mapping in soybean. The objectives of this work were to determine the location of recombination hotspots in soybean, identify genomic features associated with hotspots, and explore if hotspots are stable across two large recombinant inbred line (RIL) populations.

## MATERIALS AND METHODS

### Plant Material

The two RIL populations used in this study have been previously described by Song et al. (2016). The first RIL population was a cross between ‘Williams 82’ (*G. max*) (Bernard and Cremeens, 1988) to ‘Essex’ (*G. max*) (Smith and Camper, 1973). The second population was a cross between Williams 82 and PI479752 (*Glycine soja* Sieb. & Zucc.). The Williams 82 by Essex population (WE) consists of 922 F_5_ derived RILs developed by single seed decent (SSD) at the University of Tennessee, Knoxville, TN. The Williams 82 x PI479752 population (WP) consists of 1,083 F_5_-derved RILs and were developed by SSD at USDA-ARS, Beltsville, MD. DNA was extracted from the single F_5_ plants that derived each RIL.

### Genotyping RILs and Linkage Map Construction

Genotyping of the two RIL populations, and their linkage map construction has been previously described by Song *et al*. (2016). In brief, the WP and WE populations were genotyped with the SoySNP50K Beadchip (Song *et al*., 2013). There are 11,922 polymorphic SNPs for the WE population and 21,478 polymorphic SNPs for the WP population (Song *et al*., 2016). Since DNA was extracted from the single plants F_5_ plants that derived the RILs, genotype calls produced distinct clusters for the two homozygous and the heterozygous alleles. This allowed for highly accurate calling of each allele. Quality control steps used were the removal of markers with >10% missing data, markers with significant segregation distortion (p<0.01), and only retaining one marker if multiple markers had identical allele segregation patterns. To construct the linkage maps, MSTMAP software was used (Wu *et al*., 2008). The distance between polymorphic SNPs was calculated using JoinMap 4.0 (Van Oojien *et al*., 2006). The pericentromeric and euchromatic regions in both populations were previously defined by Song *et al*. (2016).

### Recombination Estimation and Hotspot Detection

Recombination hotspots were detected using three different methods; spline model in MareyMap (Rezvoy *et al*., 2007), a frequency metric (the ratio of the statistical genetic map to physical map distances) (Petes, 1991), and a pedigree/linkage disequilibrium (LD) method using PHASE (Li and Stephens, 2004; Rezvoy *et al*., 2007). Only MareyMap results are reported here since all three methods had comparable results (data not included). The cubic spline function in MareyMap, MMSpline 3 was used with default settings. A cubic spline consciously interpolates through second derivatives and passes through all the data points. A customized perl script was used to identify peaks (recombination hotspots) within the recombination estimate windows.

Uneven marker spacing (particularly in heterochromatic regions) led to right skewed distribution in identifying hotspot size. Therefore, the median and average differ drastically. Therefore, outliers in hotspot size distributions were identified in the data set within euchromatic and heterochromatic regions using the Tukey method (Tukey, 1977). Hotspot distribution values that were more than 1.5 times the interquartile range above the third quartile were removed from the data set.

### Correlations & Motif Discovery

To test if genomic features are associated with recombination hotspots, statistical tests for correlations were based on binomial logistical regression. The statistical tests for logistical regression were performed with Students t-test on the covariate effect using the GLM function in R 3.2.2. Soybean transposable elements were downloaded from SoyTE database, www.soybase.org (Du *et al*., 2010). The MEME suite 5.1.0 software was used to discover motifs associated with nucleotides surrounding and including the recombination hotspots within 200 b.p. upstream (Bailey *et al*., 2006). For the motif discovery, a 1st order Markov model was selected to look at both nucleotide and dinucleotide repeats across the genome and to search for motifs on both strands (Bailey *et al*., 2006). Sequence logos for each discovered motif as well as E-values were generated. Additionally, for each generated sequence motif a gene ontology analysis was completed resulting in the top five specific predictions (Buske *et al*., 2010). Also, a SpaMO analysis was performed in conjunction with the MEME suite pipeline, generating reports using the Yeast and Arabidopsis databases (Buske *et al*., 2010).

## RESULTS

The SoySNP50K Beadchip provided a high density of polymorphic markers in each of the two large RIL populations. The SoySNP50K allowed genotyping of 11,922 high quality SNPs in the WE population and 21,478 high quality SNPs in the WP population. Of the 11,922 markers in the WE population, 9,514 SNPs were located in euchromatic DNA resulting with one polymorphic marker approximately every 47 kb. The remaining 2,408 markers were located in the heterochromatic DNA resulting in an average of one polymorphic marker every 208 kb. The distribution of SNPs in the WP population between euchromatic and heterochromatic regions were similar with 17,955 markers in the euchromatic regions and 3,523 markers in the heterochromatic regions. The higher density of markers gave an average of approximately one marker per 25 kb for euchromatic regions and one marker per 142 kb for heterochromatic regions. This distribution closely resembles the original design of the SoySNP50k which has 76% of its SNPs located in the euchromatic regions and 24% of its SNPs located in heterochromatic regions.

The high density of markers genotyped with the SoySNP50K Beadchip enabled high-resolution identification of 451 recombination hotspots across both populations. The WE population contains 245 hotspots across the genome while the WP population has 206 recombination hotspots across the genome (Supplemental Table 1). Of the 451 hotspots identified, the majority (73%), are located within euchromatic DNA. Despite heterochromatic regions having severely reduced recombination (Schmutz *et al*., 2010), there were 122 hotspots identified within the heterochromatic regions of the chromosomes. To determine if these hotspots are shared between the two populations, the locations of the hotspots were also compared between the WE and WP populations. Of the 245 WE hotspots and 206 WP hotspots less than 8% occur in the same genomic locations (Supplemental Figures 1-20). This indicates that hotspot locations might not be widely conserved in bi-parental populations despite both these populations sharing a common parent.

**Figure 1.**
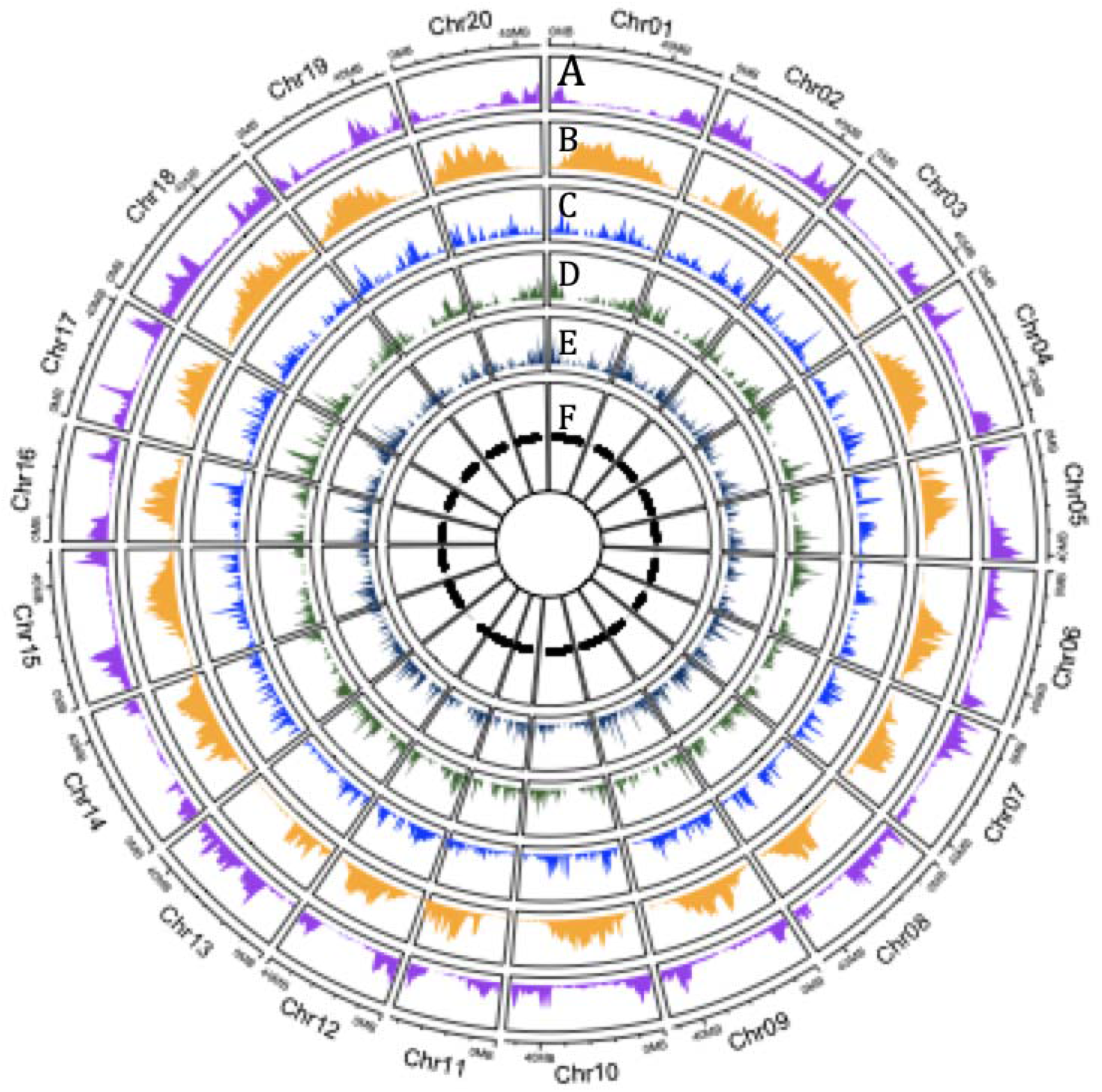
Genome wide recombination rates in two biparental populations and transposable elements associations. The two outer rings represent the average biparental populations by physical distance along chromosomes for Williams 82 x PI479752 and Williams 82 x Essex. A) directly under the physical distance ring purple displays the recombination rates in cM/Mbp (y-axis). B) The next circle in orange represents a density plot of retrotransposon (class I). C) TE Class type II MITE/Tourists element frequencies are in blue. D) TE Class type II the mite stowaway elements frequencies are in green. E) The inner circle represents heterochromatic regions in black and euchromatic regions in white

Plotting all 451 hotspots by chromosome revealed recombination rates were highest towards the distal regions of the chromosomes in euchromatic DNA and lowest in the pericentromeric regions near the centromere which are heterochromatic DNA (Figure 1). The range of recombination rate within the hotspots for both populations ranged from 0.01-20.82 cM/Mbp.

Due to the significant difference in recombination between euchromatic DNA and heterochromatic DNA, the data was split based on chromatin state (euchromatic or heterochromatic regions) and analyzed independently. Hotspot intensity for WE averaged 6.08 cM/Mbp in euchromatic DNA with a similar rate within the WP population at 6.47 cM/Mbp. The recombination rate of the two populations was not significantly different from each other (Tukey HSD; *p* =0.356). In heterochromatic pericentromeric regions of the chromosomes, hotspots were identified despite these regions having overall suppressed recombination. (Supplemental 1-20).

One example is the pericentromeric region on Chromosome 19, which contains two hotspot regions ∼329 kb apart with an average recombination rate of 16 cM/Mbp (Supplemental 19). Despite this one pericentromeric hotspot having a high recombination rate, the overall pericentromeric recombination rate was much lower, averaging 1.5 cM/Mbp across both populations. The WE population had a recombination rate of 0.89 cM/Mbp for hotspots located in pericentromeric DNA while the WP population had a higher rate of 2.1 cM/Mbp. While hotspots do occur in the highly heterochromatic pericentromeric DNA, the average strength (cM/Mbp) of the hotspot is significantly reduced by 4.78 cM/Mbp averaged across both populations compared to hotspots that occur in the euchromatic regions of the chromosomes (Tukey HSD <2e-16).

The average size of the recombination hotspots was significantly smaller in euchromatic DNA in both populations when compared to heterochromatic regions (Tukey HSD <2e-16). The average hotspot size in euchromatic regions spans 190 kb while in heterochromatic regions the average is 493 kb. Both distributions for hotspot size are skewed to the smaller size (Figure 2). This skewedness results in a median hotspot size of 85kb for euchromatic DNA and 258 kb for heterochromatic DNA. In addition, 53% of the euchromatic hotspots are less than 75 kb in length and range from 3.44 kb up to 4,317 kb (Figure 2). Heterochromatic hotspots have a range in size from 5.86 kb to 8,966 kb (Figure 2).

**Figure 2.**
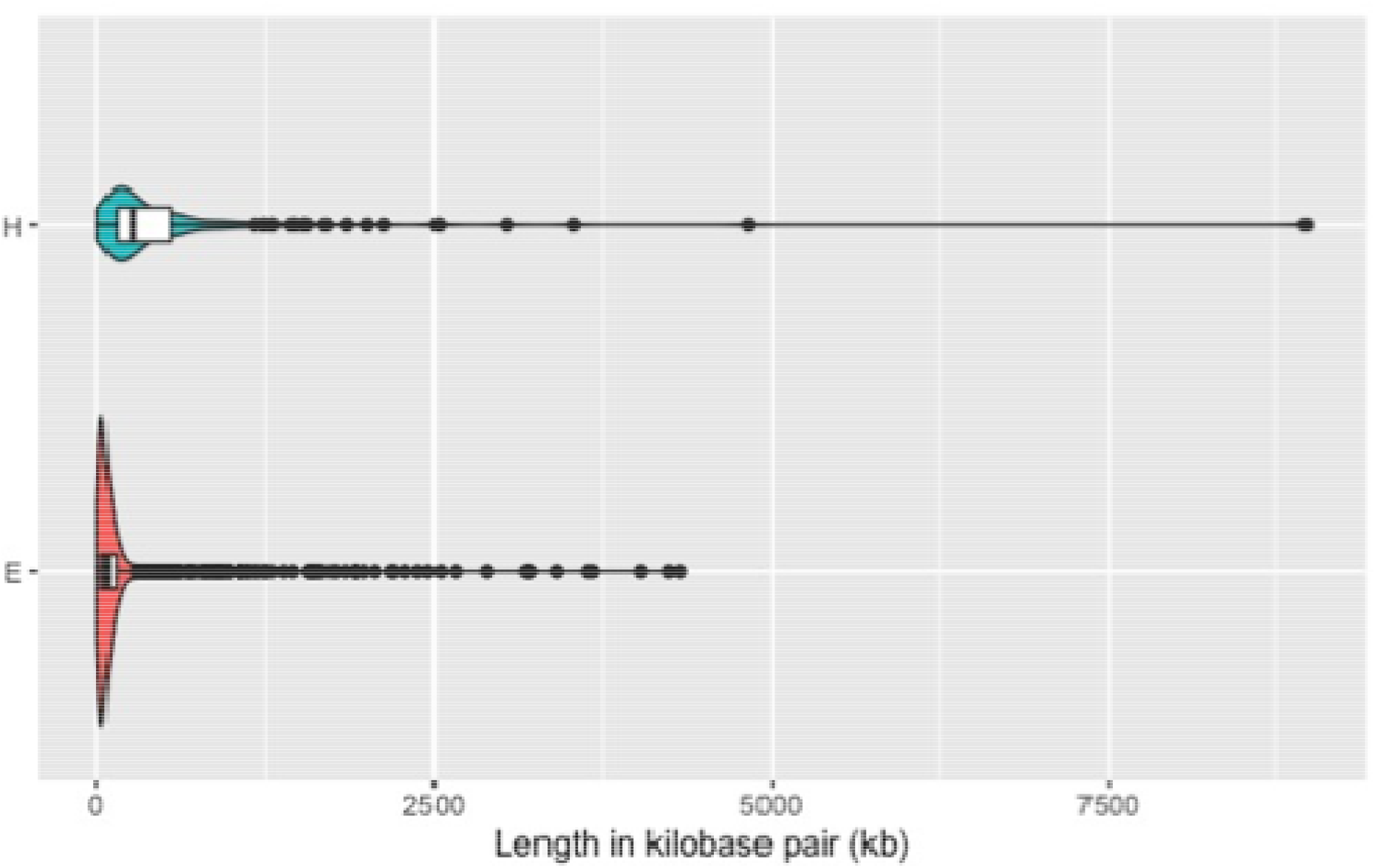
Density of hotspot length in kilobase pairs (kb) by chromatin state, heterochromatic (H) or euchromatic (E) regions. Kernel density estimates of hotspot length are represented by the plot outline. Box plot information is embedded within the violin plot. A vertical solid black line within the box represents the median and whiskers as the black horizontal line

With the location and size of each hotspot determined on a high-resolution scale, it was possible to make associations of these hotspots with genome features such as gene regions. The gene regions consist of six categories: 3’ untranslated region (UTR), 3’UTR/coding sequence (CDS), 5’UTR, 5’UTR/CDS, CDS and introns. The gene regions contained ∼28% of the identified hotspots. Of the 28% hotspots in gene regions, half were associated with introns (50.4%). The other half were distributed among the other categories with 27.9% within CDS, 13.2% within 5’UTR, 6.9% within 3’UTR, and 1.6% within 5’UTR/CDS regions.

Additional genomic features such as class I and class II transposable elements, order, super family, and other genomic descriptions from the soybean transposable element database (SoyTEeb) were tested for significant association with recombination hotspots and cold spots. Recombination hotspots were positively associated with a class of type II transposable elements, MITE/Tourist transposable elements (p-value <0.01) (Figure 3). Cold spots regions were positively associated with the long terminal repeat (LTR) order of class I transposable elements (p-value <0.005).

**Figure 3.**
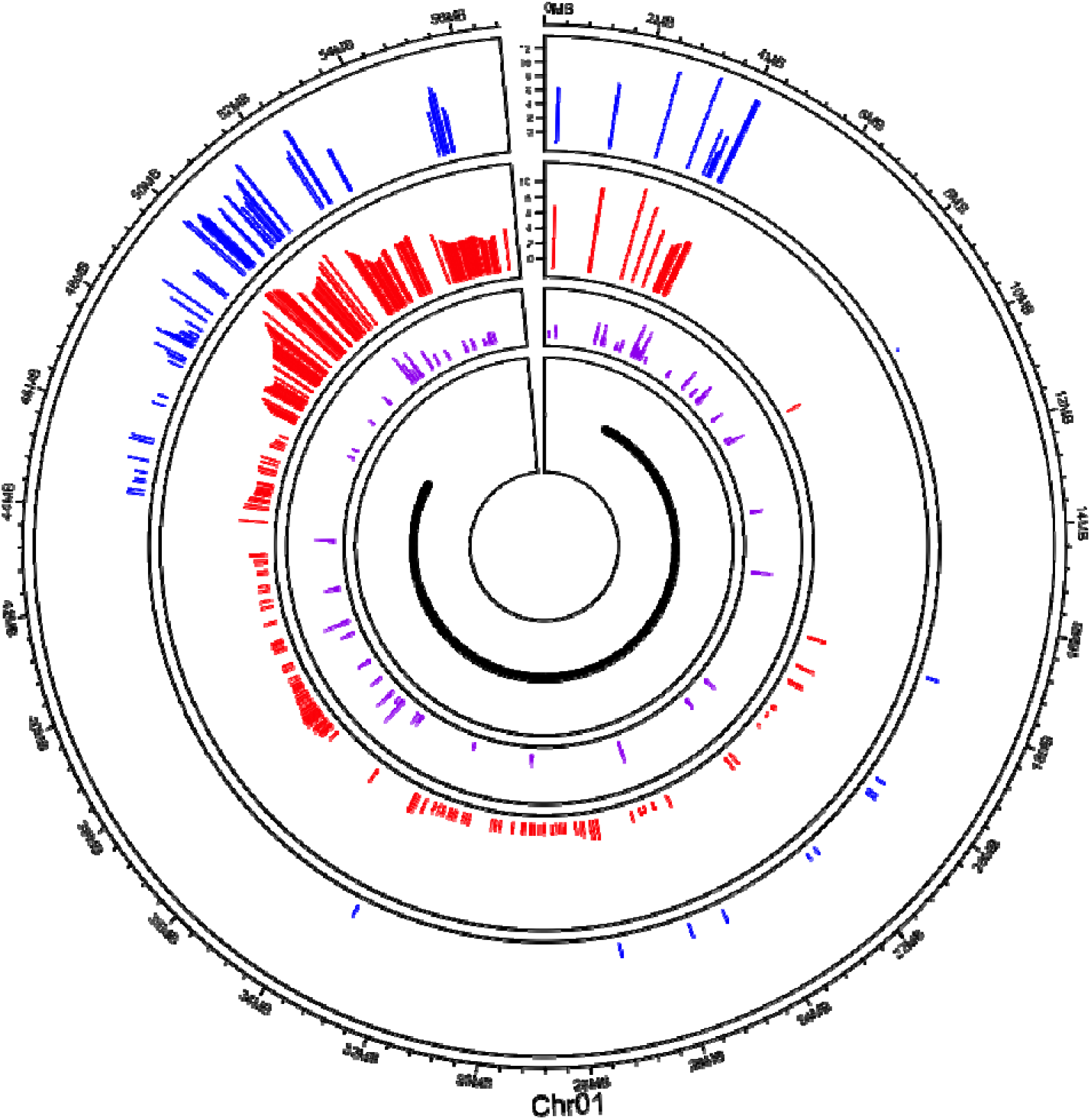
Chromosome 1 recombination rates, MITE/Tourists and heterochromatic regions. Williams 82 x PI479752 (blue) and Williams 82 x Essex (red) recombination rates are displayed in cM/Mbp (y-axis). The type II transposons subclass, MITE/Tourist are shown in purple. The black dots on the inner circle represent heterochromatic regions of the chromosome. Euchromatin regions are displayed by the grey dashed lines

Recombination associated motifs have been consistently reported in previous hotspot mapping studies (Choi and Henderson, 2015; Darrier *et al*., 2017; Marand *et al*., 2017; Mercier *et al*., 2015b). To determine if motifs are associated with recombination rates in soybean, MEME suite was used to search for over-represented motifs in the 200 bp of sequence flanking the hotspot regions. MEME reports an E value which is the estimate of the expected number of motifs with a given log likelihood ratio with the same width and site count as one would find in a similar size of random sequences. Two motifs were identified as having an association with hotspots across the genome, a poly-A motif stretch (E value = 6.8e-30) and a CCN repeat (E value = 1.7e-65) (Figure 4). The A stretch motif is associated with 32% of the hotspots in soybean. These hotspots have an average hotspot recombination rate of 7.24 cm/Mbp with the majority located in euchromatic DNA (81.8%). The hotspot intensity associated with the poly-A motif (7.24 cM/Mbp) is significantly different to hotspot regions not associated with the poly-A motif (5.84 cM/Mbp, p-value < 2.2e-16). The second motif, CNCCNCCACAACCAANNCANNA is similar to the CCN repeat that has been associated with H3K4me3 modified nucleosome (Shilo *et al*., 2015). This CCN like motif is associated with 54% of the hotspots in soybean with the majority in euchromatic DNA (78.8%). The average hotspot recombination rate associated with the CCN like motif was significantly higher (7.11 cM/Mbp) than the average recombination rate for hotspots in the two populations (WE euchromatic regions average = 6.08 cM/Mbp, WP euchromatic regions average = 6.47 cM/Mbp) (p-value = 8.633e-05). Ten percent of the hotspots were found to be associated with both motifs. Having both motifs present in a hotspot only increased the recombination rate to 7.44 cM/Mbp which was not significantly different when only one of the motifs was present in a hotspot. (p-value = 0.5281).

**Figure 4.**
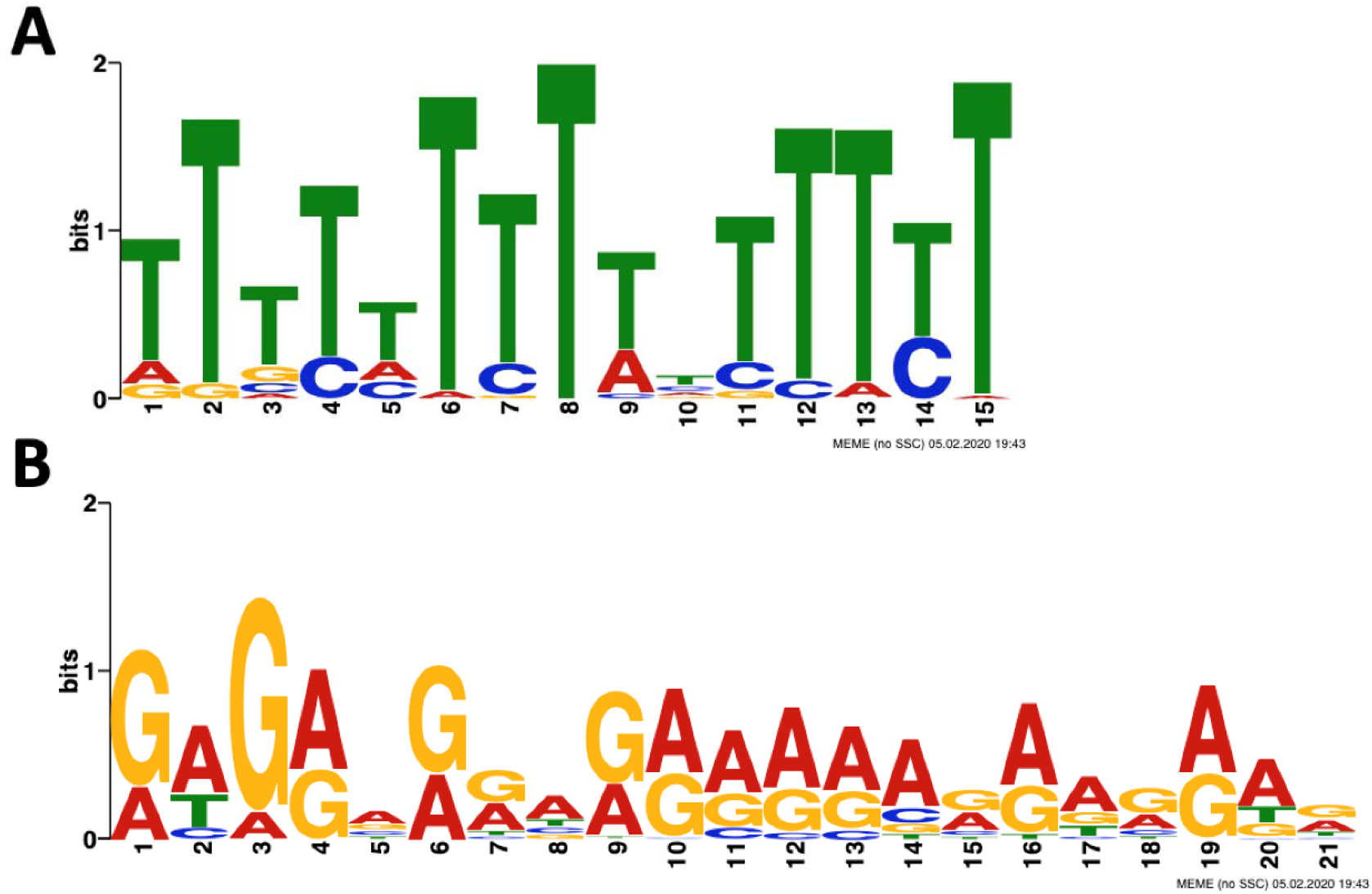
Two most common discovered motifs on recombination hotspots using MEME Suite. The Poly-T/A repeat (A) and a second motif which is CCN-like (B) were detected within 200 b.p. of recombination hotspots

A spaced motif analysis was conducted and located an overlapping sequence with the Cha4p protein for the hotspot that contains the CCN like motif. This protein encodes DNA binding transcriptional activator and mediates serine/threonine activation of the catabolic L-serine deaminase (CHA1), Zinc-finger protein Zn[2]-Cys[6] fungal type binuclear. Additionally, the gene ontology results illustrate an ∼83% association with transcriptional factor activity which is consistent with other species. In humans, Zinc-finger proteins are the largest class of transcription factors, however, they are not as common in plants. The CCCH Zinc-Finger proteins have been associated with abiotic stress tolerance (Han *et al*., 2021).

## DISCUSSION

High-density SNP data previously used to create high resolution genetic maps for two large soybean RIL populations was successfully used to map and characterize recombination hotspots in soybean. The recombination hotspot average length of 25 kb is twice as long as the 5-10 kb hotspots lengths reported in the majority of other species (Choi and Henderson, 2015; Darrier *et al*., 2017). While in horses, the recombination hotspot average size of 23.8 kb is similar to soybean (Beeson *et al*., 2019). The hotspots detected in soybean heterochromatic DNA had lengths that spanned much larger distances. The 136 kb average length of hotspots in soybean are 128 kb larger than the 8 kb average length reported in Arabidopsis heterochromatic (Drouaud *et al*., 2013a). This could be due to the hotspots in Arabidopsis being less influenced by DNA methylation (Melamed-Bessudo and Levy, 2012).

The heterochromatic DNA in soybean has been shown to have suppressed recombination (Schmutz *et al*., 2010). This led to the surprising result that 27% of the recombination hotspots discovered in the two soybean populations were discovered in heterochromatic DNA. The recombination fractions for the hotspots in the heterochromatic regions were lower than for the hotspots in euchromatic regions. However, there were exceptions. Chromosome 19 had a recombination fraction of 16 cM/Mbp, which is comparable to the euchromatic regions. The models for meiotic recombination are not fully understood or the exact role of chromatin in recombination. Recently, it was demonstrated that histone mutation can increase recombination in pericentromeric regions (Underwood *et al*., 2018). The presence of hotspots in recombination-poor regions provides future possibilities of breaking unfavorable alleles that tend to be inherited together in the large pericentromeric regions of the soybean genome.

Between the two biparental populations, less than 8% of the hotspots were located in the same genomic location. The lack of shared hotspots could be due to biased gene conversion. The potential effects of biased gene conversion have been noted by the lack of hotspots between humans and chimpanzees, despite a 99% genetic sequence similarity (Coop and Myers, 2007).

The authors discovered biased gene conversion to be a strong force in rapid decline of intense hotspots; however, dim hotspots, reduction in cM/Mbp compared to the average recombination for a genomic region, are more likely to be shared (Coop and Myers, 2007). Dim hotspots can arise by the competition between local and ancestral recombination meaning the intensity (cM/Mbp) would be suppressed, thus reducing selection against alleles causing them, as well as some alleles to rise in frequency due to genetic drift (Coop and Myers, 2007). The turnover rate of recombination hotspots was further investigated in F_1_ hybrids coming from four subspecies of *Mus musculus* with different Prdm9 alleles (Smagulova *et al*., 2016). They found preferential use of Prdm9 alleles as well as hotspots that become active in hybrids have a greater sequence diversity (Smagulova *et al*., 2016). Structural variance can influence the location of recombination hotspots along chromosomes. For example, in maize, hotspots were suppressed in regions believed to have an inversion (Rodgers-Melnick *et al*., 2015). Since structural variance in soybean is relatively low; gene content variation likely doesn’t explain the lack of hotspots shared between the two populations (McHale *et al*., 2012).

Two DNA motifs were discovered in association with recombination hotspots in soybean. One associated DNA motif was a poly-A motif and the second a CCN like repeat. The poly-A motif has been previously reported to be associated with recombination hotspots in humans, wheat, Drosophila, and Arabidopsis (Comeron *et al*., 2012; Darrier *et al*., 2017; Myers *et al*., 2008; Shilo *et al*., 2015). In wheat, the poly-A motif was found to be associated with 51.6% of the crossovers, which was the highest percentage of the reported motifs (Darrier *et al*., 2017). Poly-A motifs are thought to influence recombination hotspots due to their association with heterochromatic regions and resilience to nucleosome folding but do not directly prompt recombination (Comeron *et al*., 2012; Darrier *et al*., 2017; Myers *et al*., 2008; Shilo *et al*., 2015). The second associated motif, a CNN-like repeat, shares similar characteristics to previously identified hotspot motifs in wheat, Arabidopsis, dog, Drosophila (Auton *et al*., 2013; Comeron *et al*., 2012; Darrier *et al*., 2017; Shilo *et al*., 2015). However, the CCN repeat motif was not

associated with recombination hotspot in maize (Rodgers-Melnick *et al*., 2015). The two motifs did have significantly higher rates of recombination when compared to hotspots without the motifs (p-value < 2.2e-16). This could indicate that the motifs play a role in increasing the probability of a recombination when compared to other hotspots without the motifs. Although the two motifs have been associated with hotspots in other species, the association of recombination intensity as a separate metric was not reported.

Short motifs do not completely explain variation in recombination hotspots; other genomic features such as transposable elements heavily influence recombinant positions. Darrier *et al*. (2017) showed the importance of transposable elements in wheat recombination hotspots by reporting retrotransposons (GYPSY and COPIA elements) are 63.7% higher in non-recombinant regions. In general, plant genomes contain a large number of retrotransposons. The soybean genome contains 42.24% retrotransposons (Schmutz *et al*., 2010). Therefore, it is not surprising that the LTR subclass (41.99% of the genome) is positivity associated with recombination cold spots (p-value <0.005).

DNA transposons (Type II) are less abundant than type I. Soybean contains only 16.5% of type II transposable elements (Schmutz *et al*., 2010). Terminal inverted repeats make up the largest order within the DNA transposons in soybean and the largest subclass belongs to CACTA (10.16%) (Schmutz *et al*., 2010). One of the smallest groups is PIF/Harbinger covering only 0.29% and the MITE tourist subclass with 0.33% of the genome (Schmutz *et al*., 2010). In soybean, recombination hotspots display significant association with and MITE Tourist elements (p-value <0.01). Potato hotspots have been associated with MITE Stowaway elements while rice hotspots have been associated with PIF/Harbinger (Darrier *et al*., 2017; Marand *et al*., 2017).

Soybean and potato are both palaeopolyploid. Wheat hotspots have been associated with TC1-mariner elements, (Marand *et al*., 2019). Marand *et al*. (2019) hypothesized an indirect role for Stowaway and PIF/Harbinger elements in promoting long AT-repeat regions over time, creating regions of DNA instability and susceptibility to double strand breaks. The TC1-Mariner elements insertion site is in a TA sequence and are found close to genic regions, which is very similar to the Tourist target repeat region, TAA (Darrier *et al*., 2017; Zhao *et al*., 2016). MITE’s capabilities to alter sequences near gene regions could attract DNA binding domains of meiotic factors, similar to PRDM9 (Myers *et al*., 2008).

## CONCLUSION

Within two large segregating soybean populations, recombination hotspots were identified and characterized. Hotspots in these populations were more commonly found in euchromatic regions with a quarter of them located in heterochromatic DNA. The small percentage of hotspots shared between the populations could allude to a role of gene content variation affecting the location of hotspots. Recombination cold spots were found to be associated with LTR transposable elements. Two common motifs, an A stretch and a CCN, were found in association with recombination hotspots that had a higher rate of recombination than hotspots without the motifs. Uniquely, soybean hotspots are found in areas enriched with Tourist MITEs, which reside in a very small percentage of the genome. While other classifications of MITEs were observed in association with hotspots for potato, rice, and wheat, each one classifies in the TIR order. Further investigation of individual MITE elements such as Tourist, Stowaway, and Tc1-mariner elements near recombination hotspots will be necessary to identify a potential mechanism.

Collectively, this study provides insights into the distinctive and shared genomic features of soybean recombination hotspots.

## ACKNOWLEDGEMENT

Funding for this project was provided by University of Nebraska Lincoln Agriculture Research Division and the University of Nebraska-Lincoln Agronomy and Horticulture Department. This work was completed utilizing the Holland Computing Center of the University of Nebraska, which receives support from the Nebraska Research Initiative.

## SUPPLEMENTAL MATERIAL

Supplemental Figures 1-20 **Recombination hotspots for each chromosome**. The two outer rings display the recombination rates in cM/Mbp (y-axis) for each biparental populations by physical distance along chromosomes, Williams 82 x PI479752 (blue) and Williams 82 x Essex (red). The inner most circle displays the significant hotspots for each population in their respective color.

Supplemental Table 1 **Recombination hotspot location, size, and recombination rate in each population**

