## Supplemental Figure for "RECOMBINATION HOTSPOTS IN SOYBEAN [*GLYCINE MAX* (L.) MERR.]"

**Supplementary Fig S1. Chromosome 1 Recombination Hotspots.**

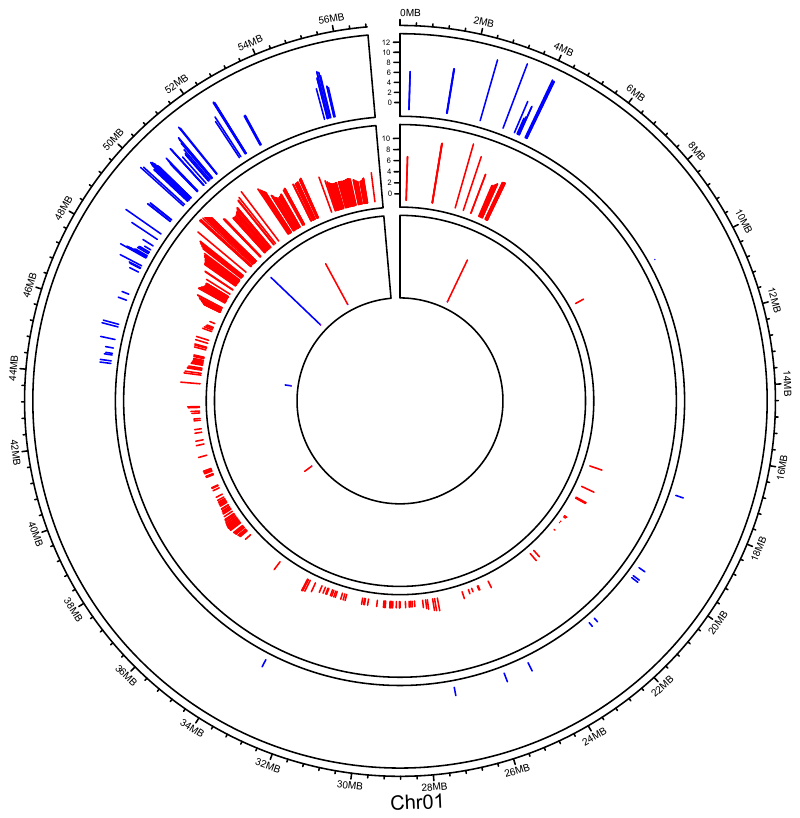

**Supplementary Fig S2. Chromosome 2 Recombination Hotspots.**

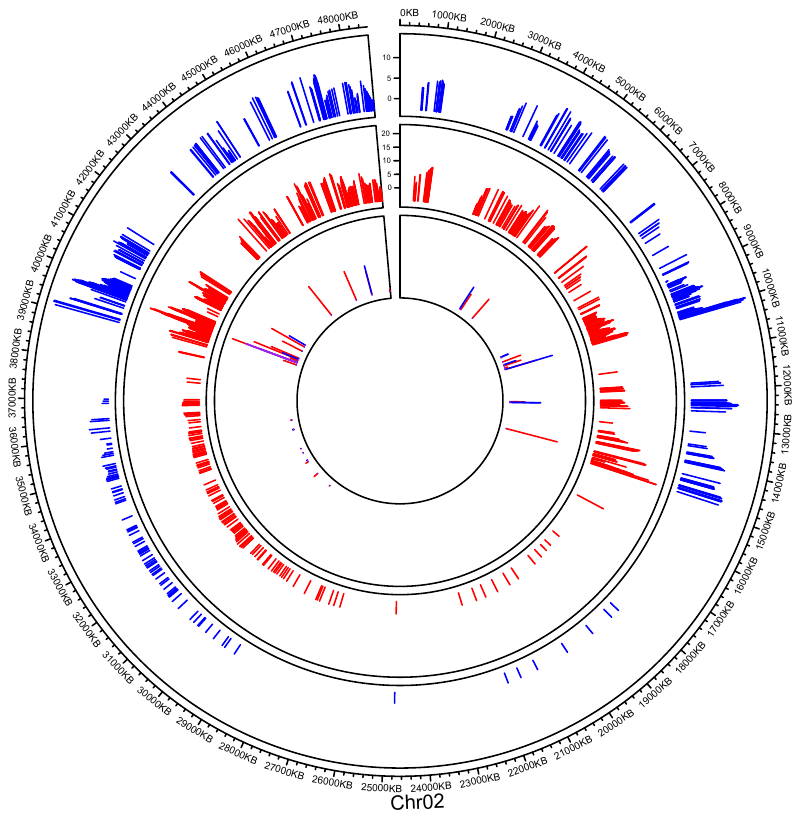

**Supplementary Fig S3. Chromosome 3 Recombination Hotspots.**

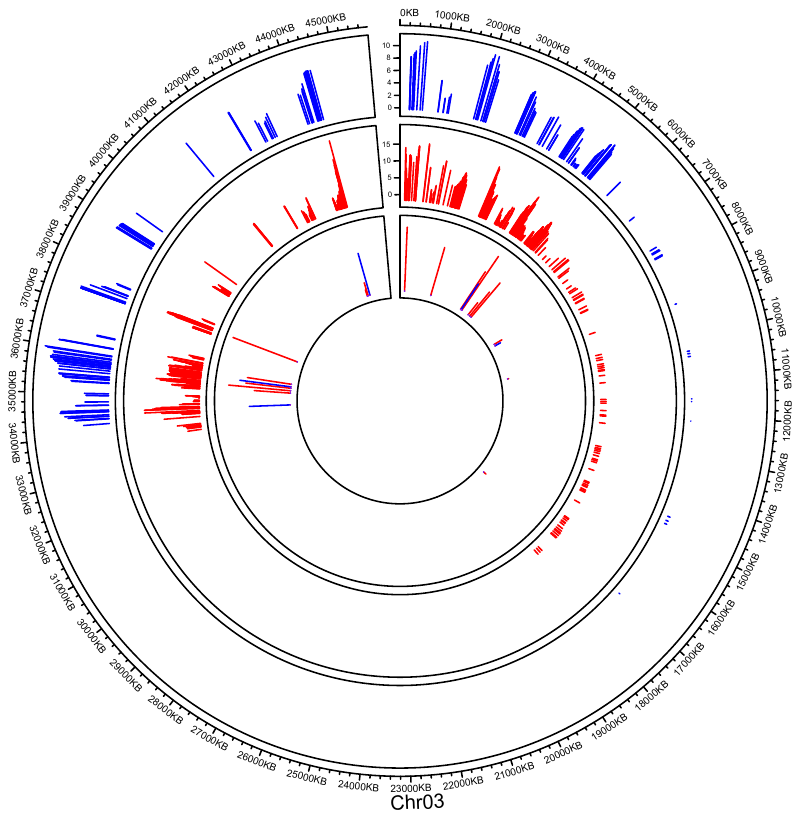

**Supplementary Fig S4. Chromosome 4 Recombination Hotspots.**

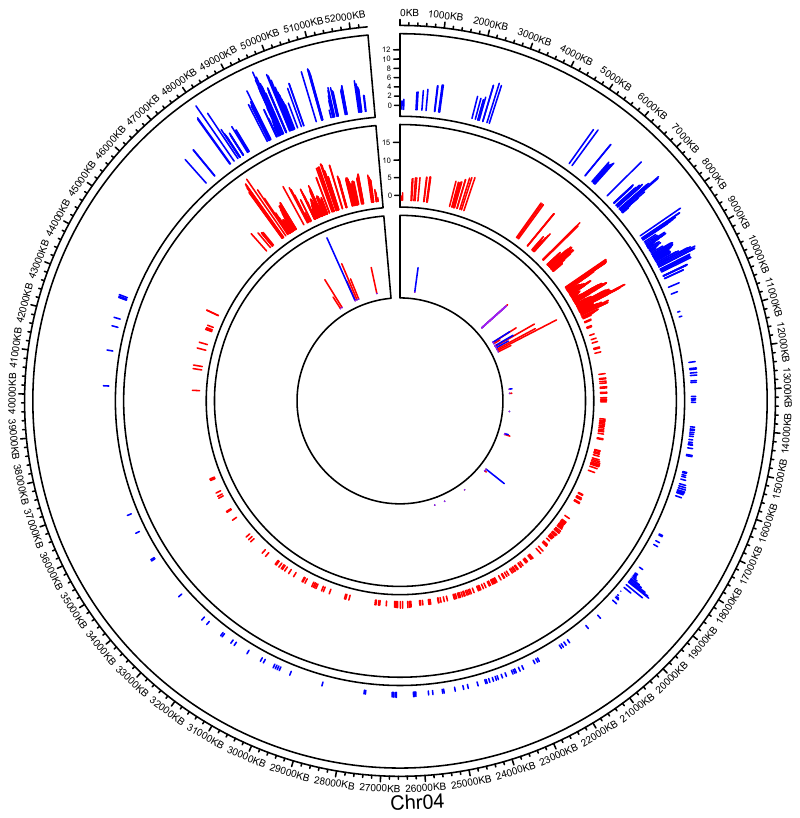

**Supplementary Fig S5. Chromosome 5 Recombination Hotspots.**

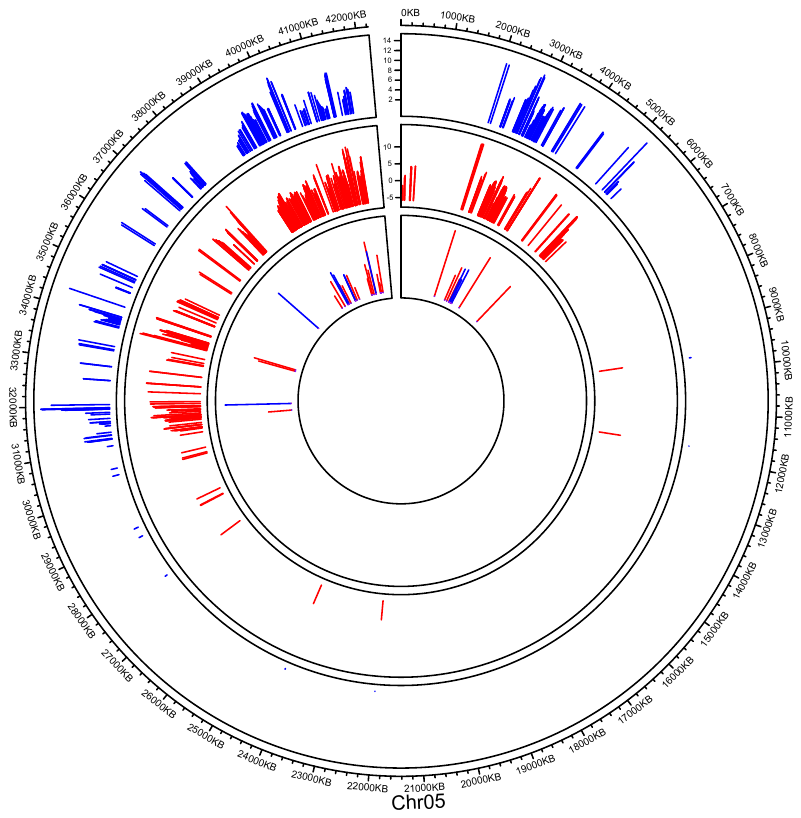

**Supplementary Fig S6. Chromosome 6 Recombination Hotspots.**

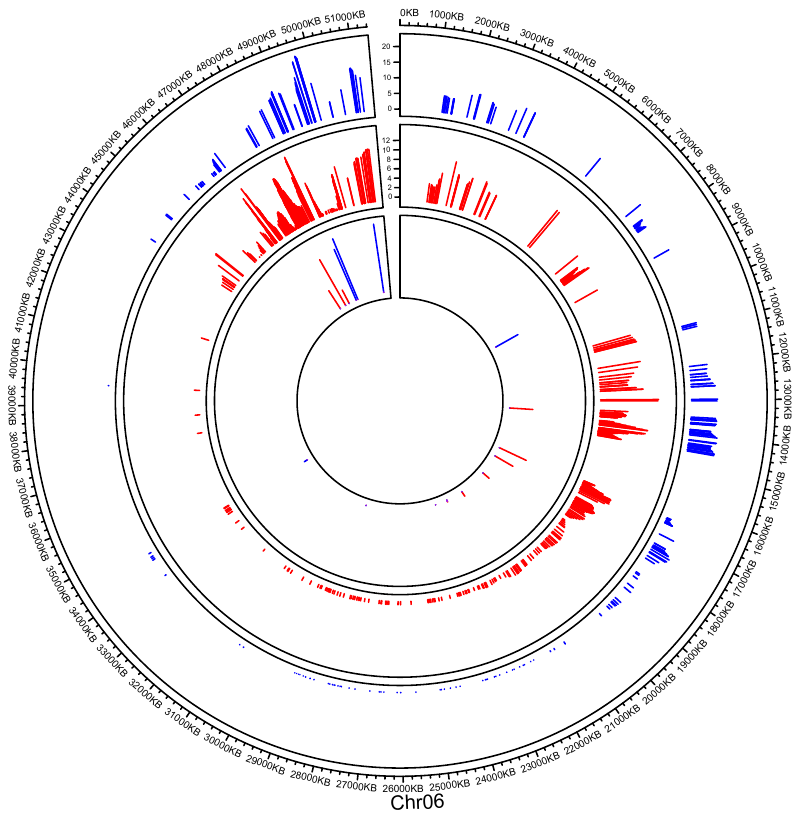

**Supplementary Fig S7. Chromosome 7 Recombination Hotspots.**

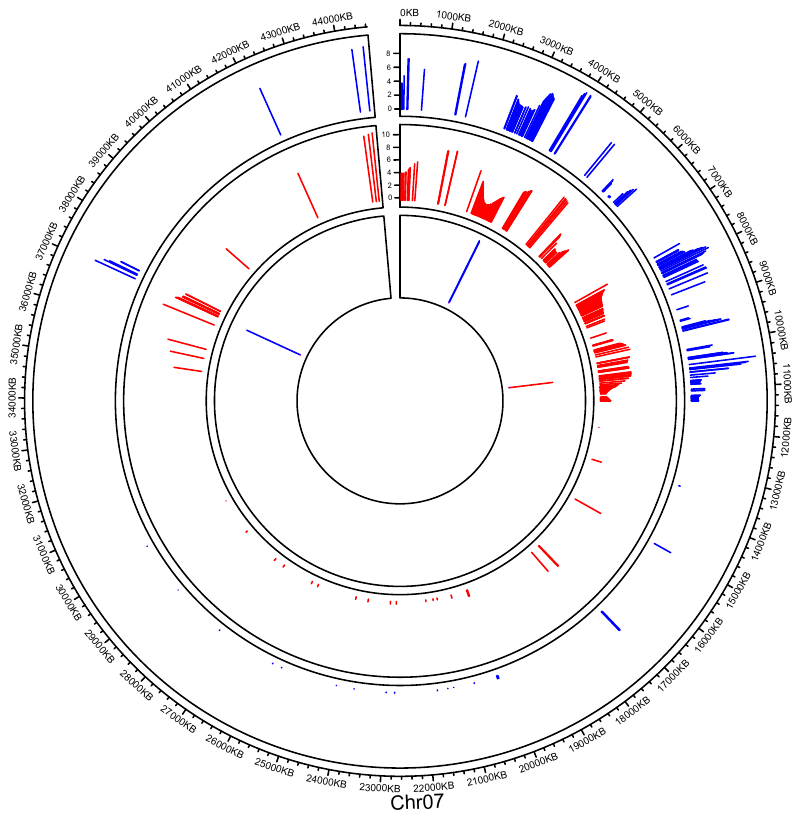

**Supplementary Fig S8. Chromosome 8 Recombination Hotspots.**
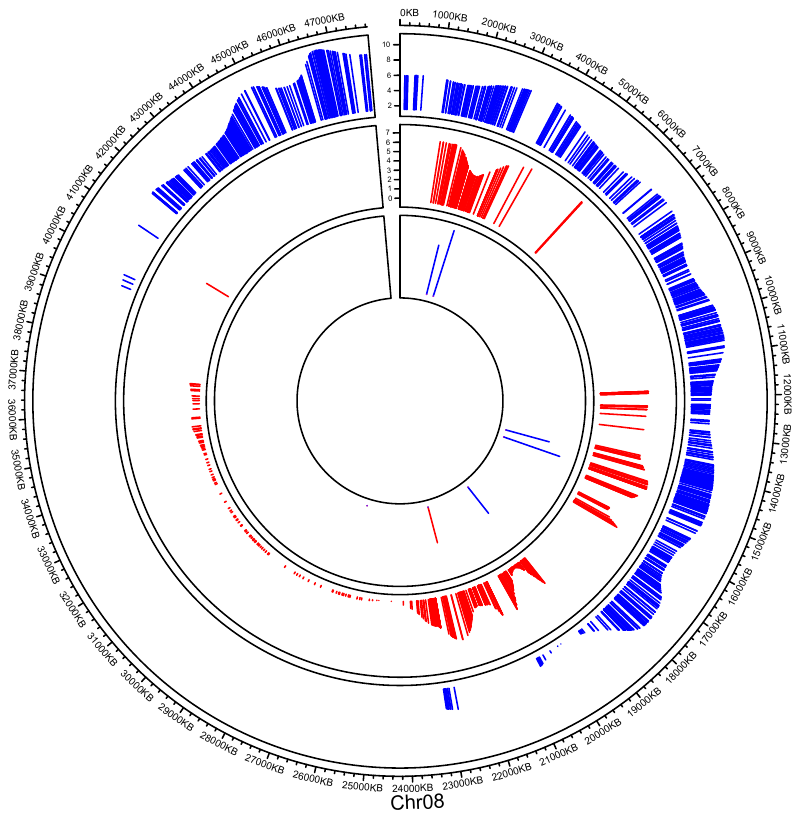

**Supplementary Fig S9. Chromosome 9 Recombination Hotspots.**

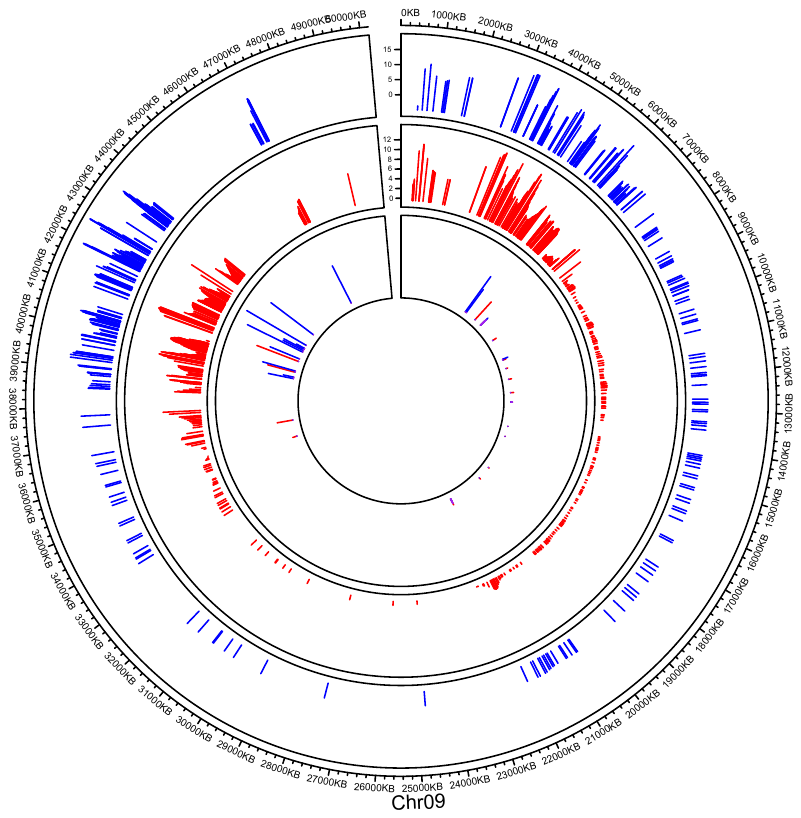

**Supplementary Fig S10. Chromosome 10 Recombination Hotspots.**

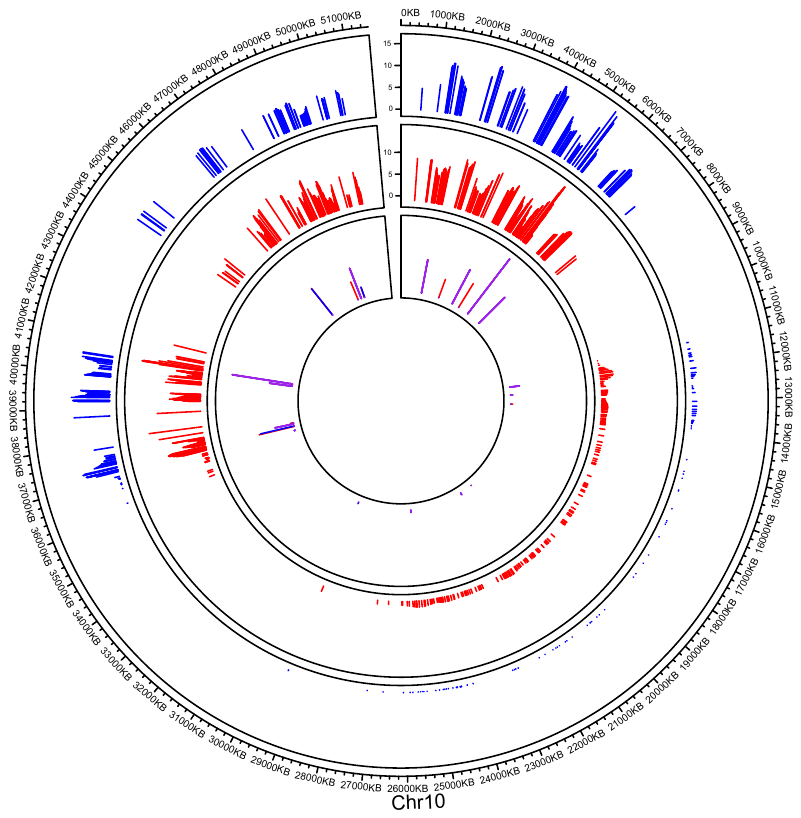

**Supplementary Fig S11. Chromosome 11 Recombination Hotspots.**

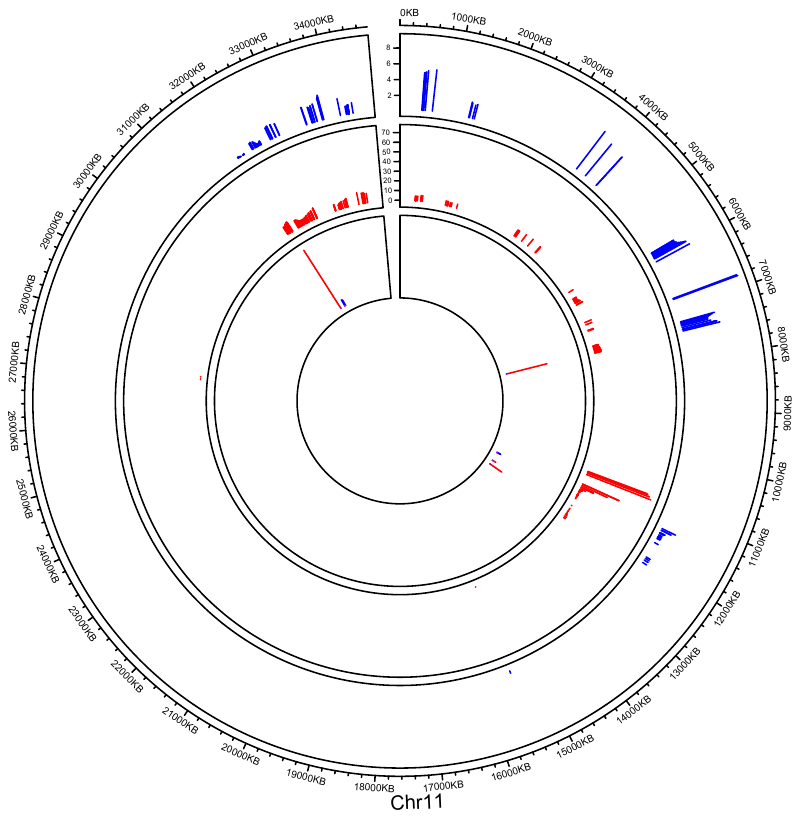

**Supplementary Fig S12. Chromosome 12 Recombination Hotspots.**

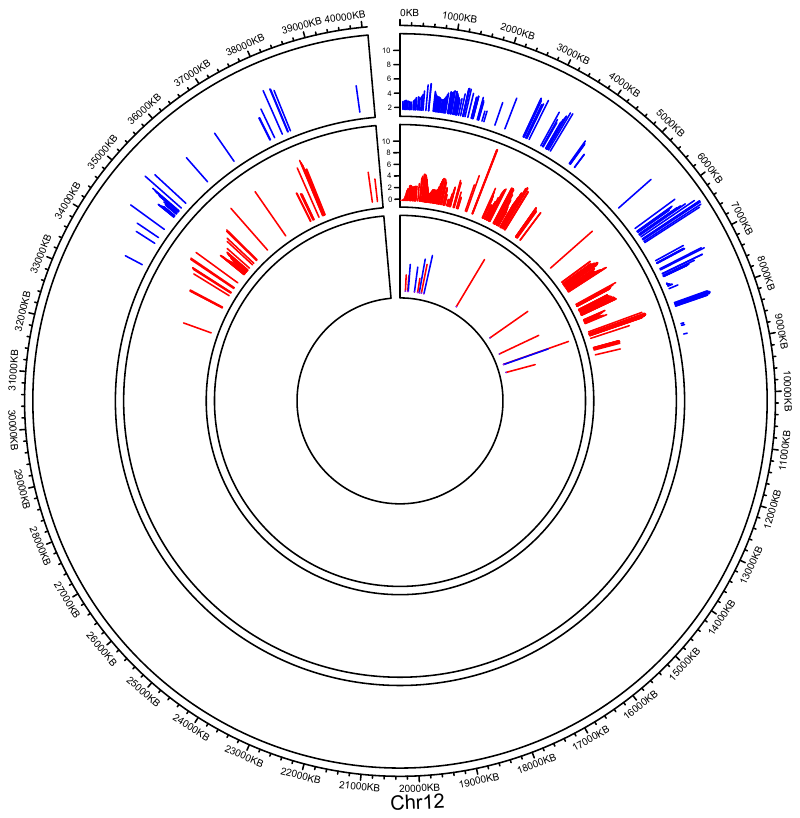

**Supplementary Fig S13. Chromosome 13 Recombination Hotspots.**

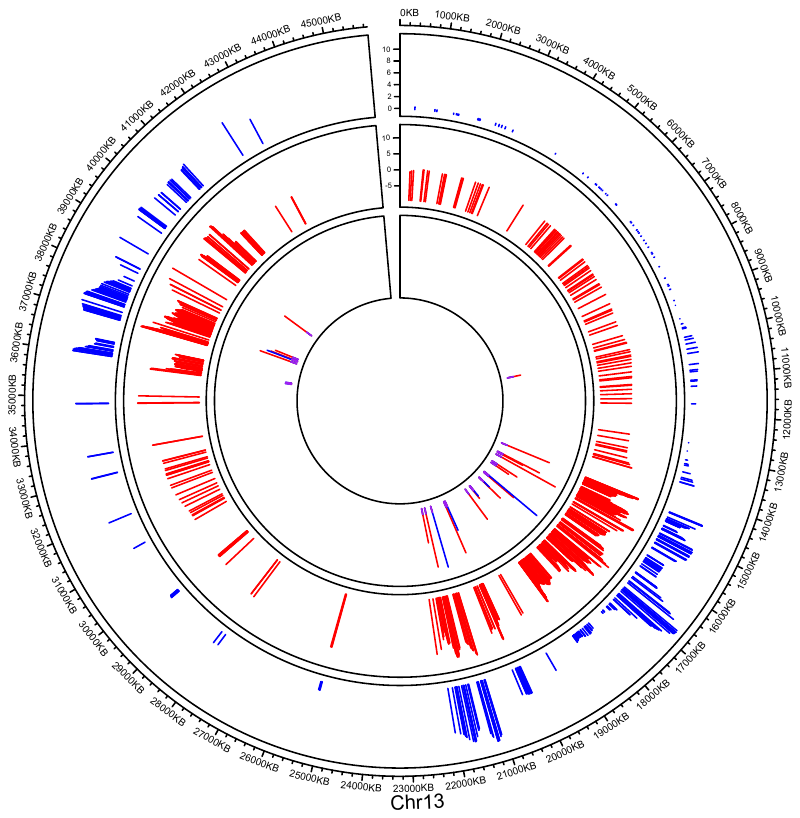

**Supplementary Fig S14. Chromosome 14 Recombination Hotspots.**

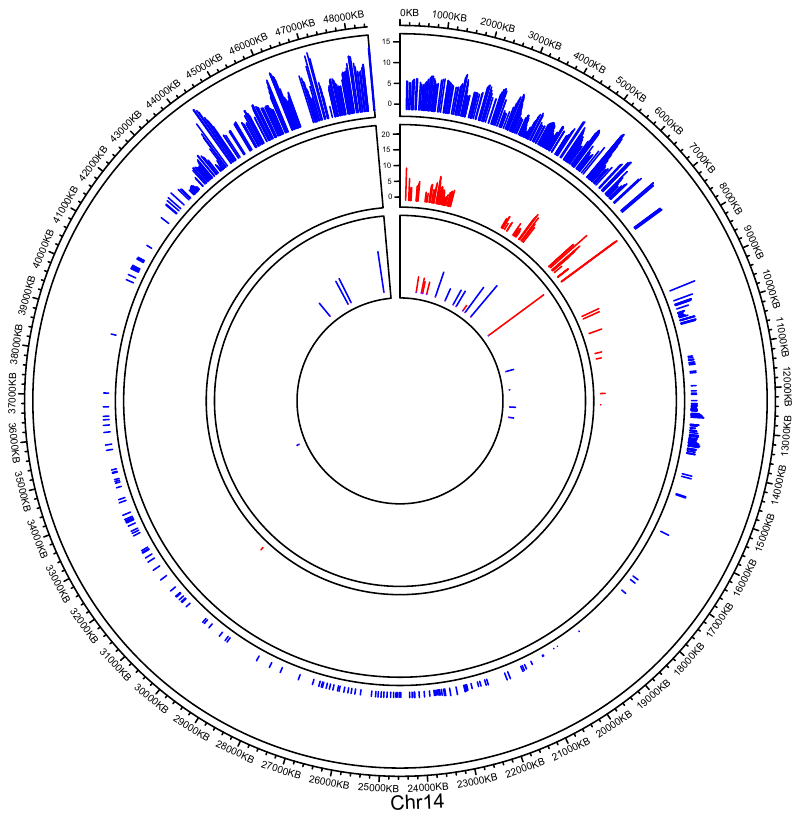

**Supplementary Fig S15. Chromosome 15 Recombination Hotspots.**

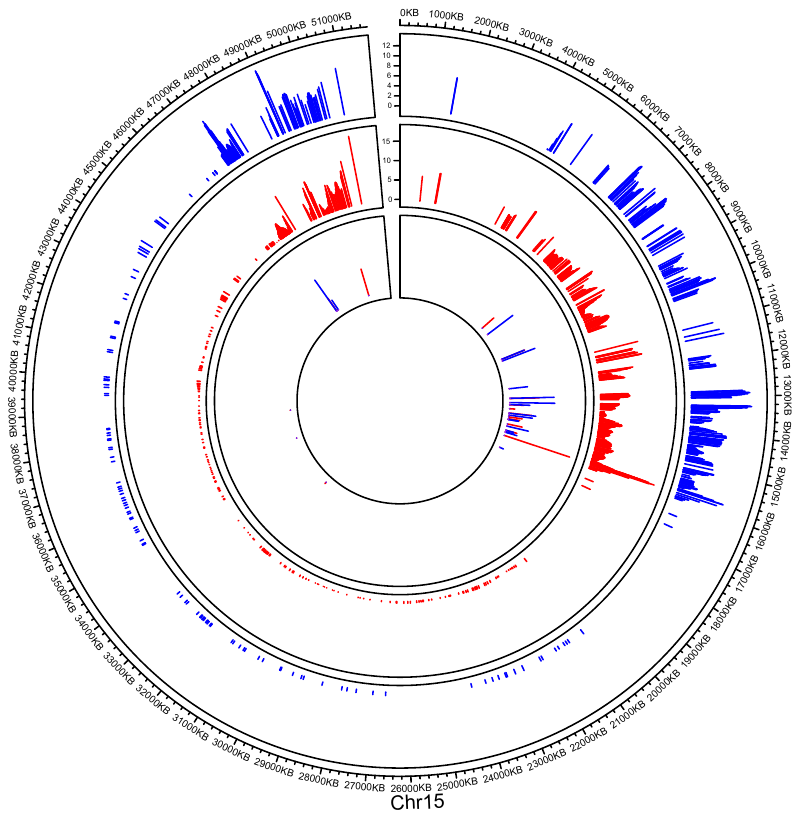

**Supplementary Fig S16. Chromosome 16 Recombination Hotspots.**

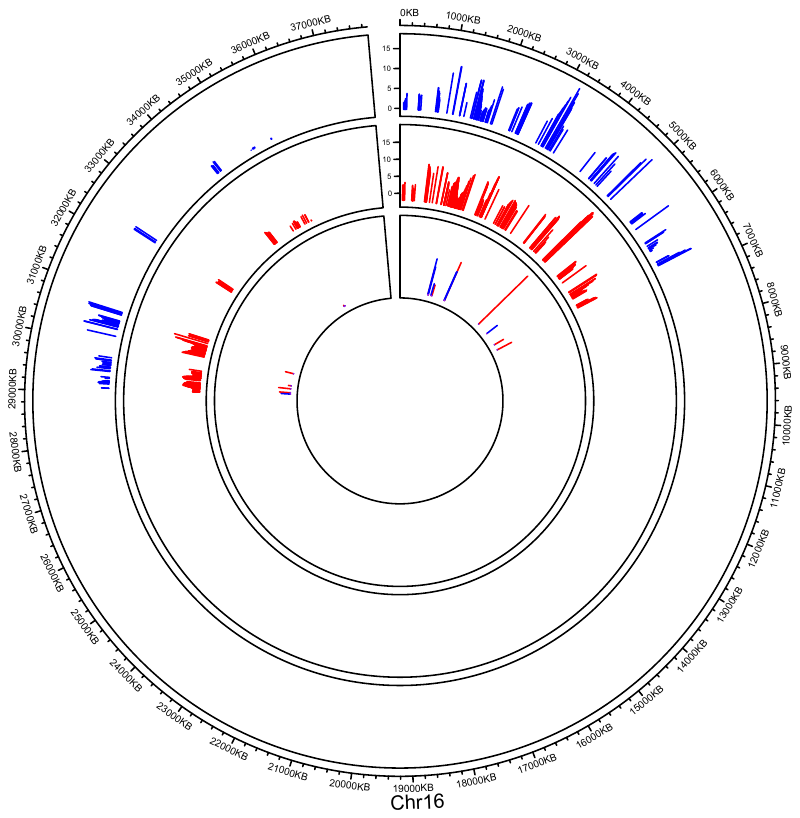

**Supplementary Fig S17. Chromosome 17 Recombination Hotspots.**

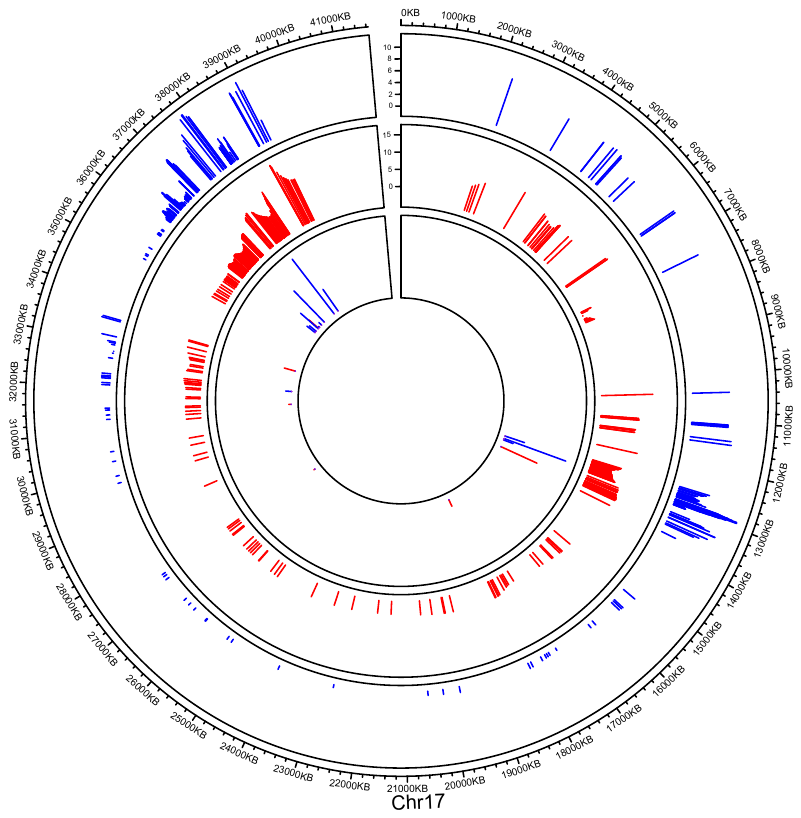

**Supplementary Fig S18. Chromosome 18 Recombination Hotspots.**

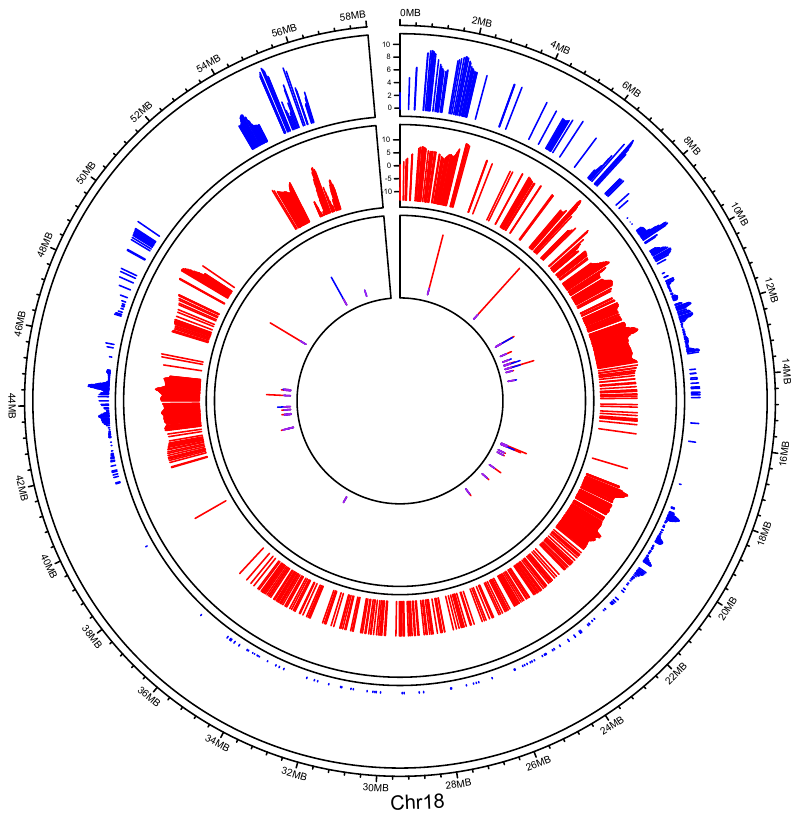

**Supplementary Fig S19. Chromosome 19 Recombination Hotspots.**

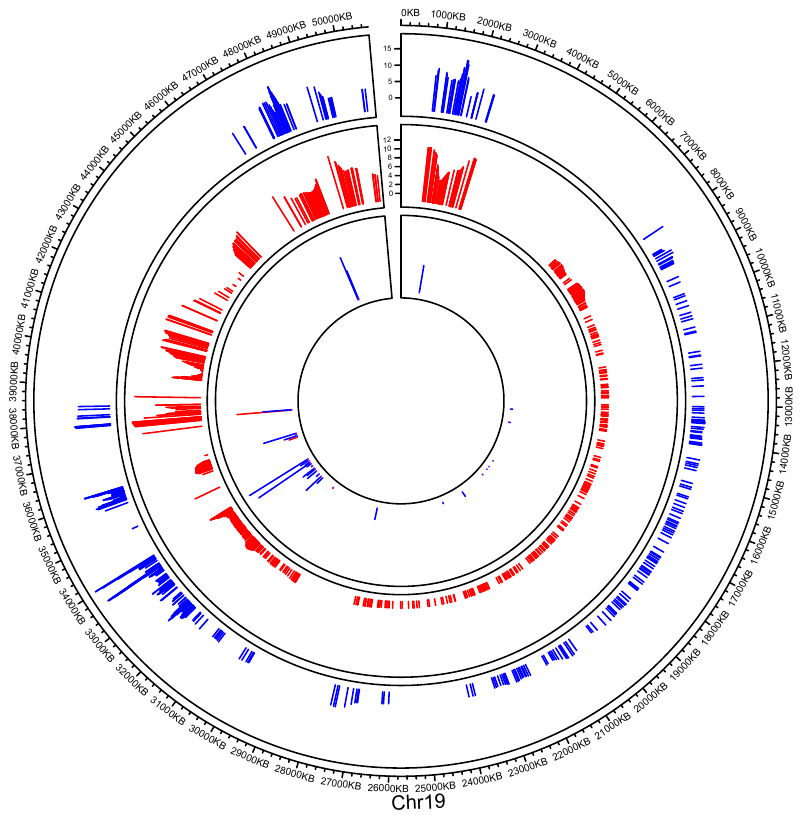

**Supplementary Fig S20. Chromosome 20 Recombination Hotspots.**

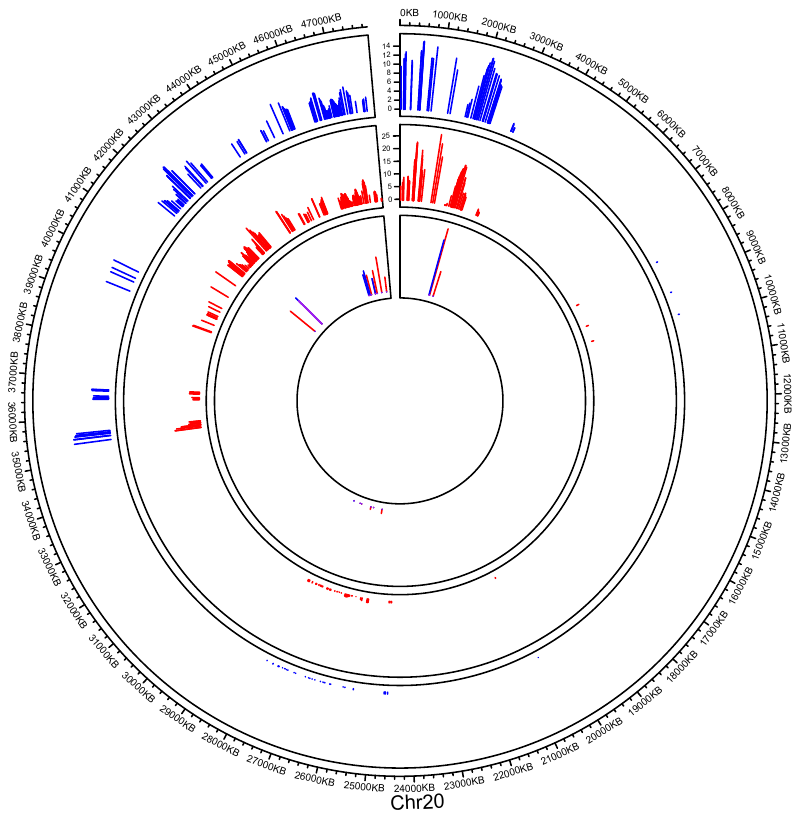
